# Rotavirus as an Expression Platform of the SARS-CoV-2 Spike Protein

**DOI:** 10.1101/2021.02.18.431835

**Authors:** Asha A. Philip, John T. Patton

## Abstract

Rotavirus, a segmented double-stranded RNA virus, is a major cause of acute gastroenteritis in young children. The introduction of live oral rotavirus vaccines has reduced the incidence of rotavirus disease in many countries. To explore the possibility of establishing a combined rotavirus-SARS-CoV-2 vaccine, we generated recombinant (r)SA11 rotaviruses with modified segment 7 RNAs that contained coding sequences for NSP3 and FLAG-tagged portions of the SARS-CoV-2 spike (S) protein. A 2A translational element was used to drive separate expression of NSP3 and the S product. rSA11 viruses were recovered that encoded the S-protein S1 fragment, N-terminal domain (NTD), receptor-binding domain (RBD), extended receptor-binding domain (ExRBD), and S2 core (CR) domain (rSA11/NSP3-fS1, -fNTD, -fRBD, -fExRBD, and -fCR, respectively). Generation of rSA11/fS1 required a foreign-sequence insertion of 2.2-kbp, the largest such insertion yet made into the rotavirus genome. Based on isopycnic centrifugation, rSA11 containing S sequences were denser than wildtype virus, confirming the capacity of the rotavirus to accommodate larger genomes. Immunoblotting showed that rSA11/-fNTD, -fRBD, -fExRBD, and -fCR viruses expressed S products of expected size, with fExRBD expressed at highest levels. These rSA11 viruses were genetically stable during serial passage. In contrast, rSA11/NSP3-fS1 failed to express its expected 80-kDa fS1 product, for unexplained reasons. Moreover, rSA11/NSP3-fS1 was genetically unstable, with variants lacking the S1 insertion appearing during serial passage. Nonetheless, these results emphasize the potential usefulness of rotavirus vaccines as expression vectors of portions of the SARS-CoV-2 S protein (e.g., NTD, RBD, ExRBD, and CR) with sizes smaller than the S1 fragment.

**Importance:** Among the vaccines administered to children in the US and many other countries are those targeting rotavirus, a segmented double-stranded RNA virus that is a major cause of severe gastroenteritis. In this study, we have examined the feasibility of modifying the rotavirus genome by reverse genetics, such that the virus could serve as an expression vector of the SARS-CoV-2 spike protein. Results were obtained showing that recombinant rotaviruses can be generated that express domains of the SARS CoV-2 spike protein, including the receptor-binding domain (RBD), a common target of neutralizing antibodies produced in individuals infected by the virus. Our findings raise the possibility of creating a combined rotavirus-COVID-19 vaccine that could be used in place of current rotavirus vaccines.

## INTRODUCTION

The impact of severe acute respiratory syndrome coronavirus 2 (SARS-CoV-2) on human mortality and morbidity has stimulated broad ranging efforts to develop vaccines preventing coronavirus disease 19 (COVID-19) (1,2). Given that the virus can cause asymptomatic and symptomatic infections in individuals of all ages, including infants and young children, comprehensive strategies to control the SARS-CoV-2 pandemic may require modification of childhood immunization programs to include COVID-19 vaccines (3,4). Among the vaccines routinely administered to infants in the US and many other countries, are those targeting rotavirus, a segmented double-stranded RNA (dsRNA) virus that is a primary cause of severe acute gastroenteritis (AGE) in children during the first 5 years of life (5). The most widely used rotavirus vaccines are given orally and formulated from live attenuated virus strains (6). These vaccines induce the production of neutralizing IgG and IgA antibodies (7,8,9) and have been highly effective in reducing the incidence of rotavirus hospitalizations and mortality (10,11).

Advances in rotavirus reverse genetics technologies have allowed the generation of recombinant rotaviruses that serve as expression platforms of heterologous proteins (12–19). The rotavirus genome consists of 11 segments of dsRNA, with a total size of ~18.6 kbp for group A strains (rotavirus species A) typically associated with pediatric AGE (20). Most of the genome segments contain a single open-reading frame (ORF); these encode the 6 structural (VP1-VP4, VP6-VP7) or 6 nonstructural (NSP) viral proteins (21). The recently-developed rotavirus reverse genetics systems consist of eleven T7 transcription (pT7) vectors, each directing synthesis of a unique viral (+)RNA when transfected into baby-hamster kidney cells producing T7 RNA polymerase (BHK-T7 cells). In some cases, support plasmids expressing capping enzymes [African swine fever virus NP868R (22) or vaccinia virus D1L/D12R (18)] or fusion proteins [avian reovirus p10FAST (18)] are co-transfected with the pT7 vectors to enhance recovery of recombinant viruses. Rotavirus reverse genetics systems have been used to mutate several of the viral genome segments and to generate virus strains that express reporter proteins (13,17,23–26).

Genome segment 7 of group A rotaviruses encodes NSP3 (36 kDa), an RNA-binding protein that acts a translation enhancer of viral (+)RNAs and is expressed at moderate levels in infected cells (27,28). In a previous study, we showed that the single NSP3 ORF could be re-engineered by reverse genetics to express two separate proteins through placement of a teschovirus 2A translational stop-restart element at the end of the NSP3 ORF, followed by the coding sequence for a heterologous protein (17). Through this approach, well-growing genetically-stable recombinant rotaviruses have been generated that express NSP3 and one or more fluorescent proteins (FPs) [e.g., mRuby (red), UnaG (green), TagBFP (blue), etc.] from segment 7, an advance allowing study of rotavirus biology by live cell imaging (15). The NSP3 product of these recombinant viruses is functional, capable of dimerization and inducing the nuclear accumulation of the cellular poly(A)-binding protein (16,17). Thus, recombinant rotaviruses that express foreign proteins via addition of a 2A element and coding sequence into segment 7 downstream of the NSP3 ORF retain the full complement of functional viral ORFs.

As a step towards developing a combined rotavirus-SARS-CoV-2 vaccine, we explored the possibility of generating recombinant rotaviruses that express regions of the SARS-CoV-2 spike (S) protein through re-engineering of the NSP3 ORF in segment 7. Trimers of the S protein form crown-like projections that emanate from the lipid envelop surrounding the SARS-CoV-2 virion (29,30). Cleavage of the trimeric spikes by extracellular furin-like proteases generates S1 and S2 fragments, each which possess activities essential for virus entry (Fig. 1). The S1 fragment includes an N-terminal domain (NTD) and a receptor-binding domain (RBD), the latter mediating virus interaction with the cell surface receptor angiotensin-converting enzyme 2 (ACE2) (31). The S2 fragment is responsible for S-protein trimerization and contains fusion domains that are essential for virus entry. SARS-CoV-2-specific antibodies with neutralizing activity have been mapped to various regions of the S protein, including the NTD, RBD, and fusion domains (32–35). We determined that by inserting S coding sequences into rotavirus genome segment 7 downstream of the NSP3 ORF and a 2A element, well-growing genetically-stable recombinant rotaviruses can be made that express all or portions of the S1 and S2 fragment. These findings raise the possibility of constructing rotavirus vaccine strains that are not only capable of inducing immunological protective responses against rotavirus, but also COVID-19.

**Figure 1.**
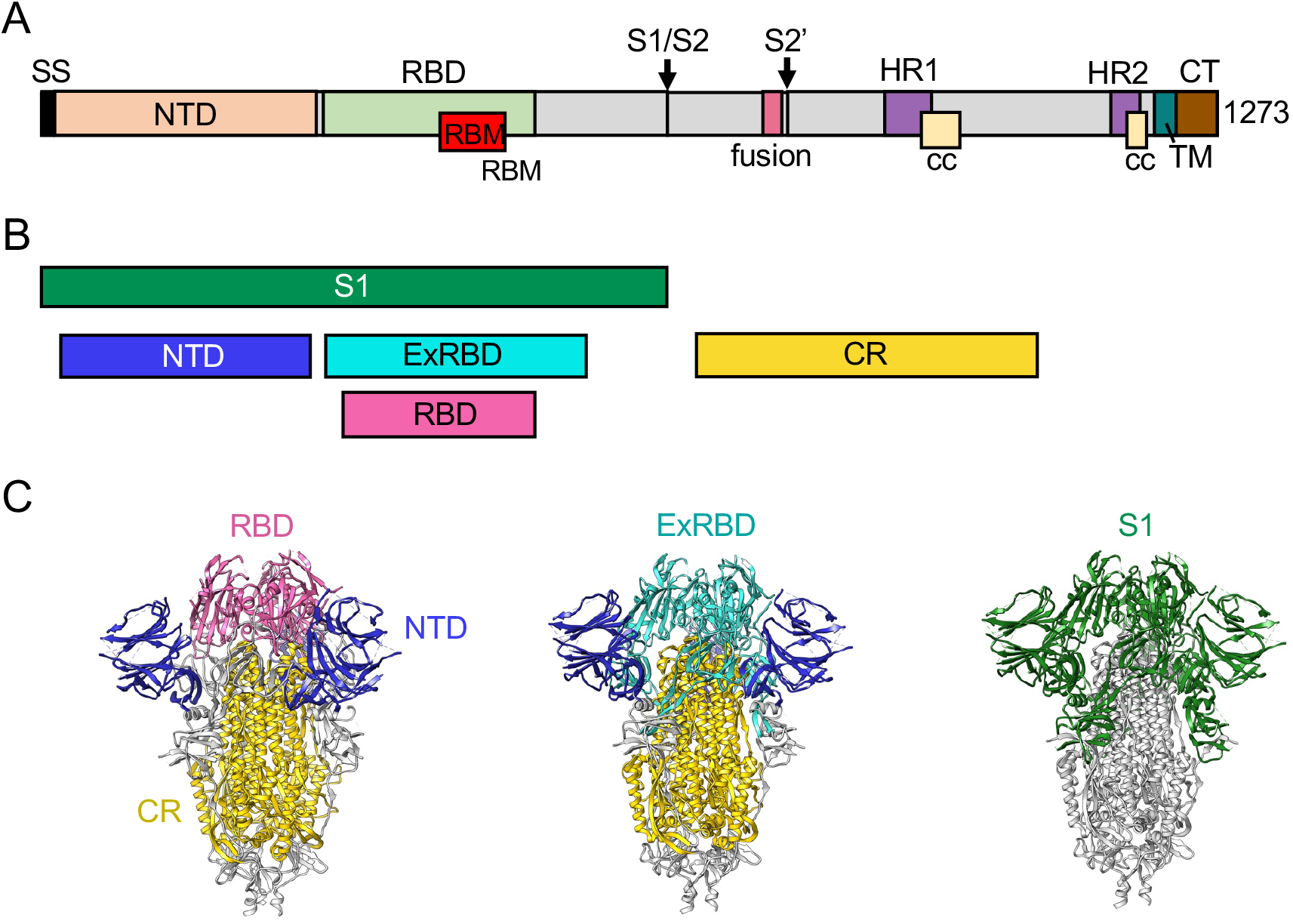
Domains of the SARS-CoV-2 S protein expressed by rSA11. **(A)** S protein trimers are cleaved at the S1/S2 junction by furin proconvertase and at the S2’ site by the TMPRSS2 serine protease. The S1 fragment contains a signal sequence (SS), N-terminal domain (NTD), receptor binding domain (RBD), and receptor binding motif (RBM). The S2 fragment contains a trimeric core region, transmembrane anchor (TM), and fusion domain. **(B)** Portions of the S protein expressed by recombinant rotaviruses are indicated. **(C)** Ribbon representations of the closed conformation of the trimeric S protein (PDB 6VXX) showing locations of the RBD (magenta), extended RBD (ExRBD, cyan), NTD (blue), core (CR, gold) domains and the S1 cleavage product (green).

## RESULTS AND DISCUSSION

### Modified segment 7 (NSP3) expression vectors containing SARS-CoV-2 S sequences

To examine the possibility of using rotavirus as an expression platform for regions of the SARS-CoV-2 S protein, we replaced the NSP3 ORF in the pT7/NSP3SA11 transcription vector with a cassette comprised of the NSP3 ORF, a porcine teschovirus 2A element, and a coding sequence of the S protein (Fig. 2). The cassette included a flexible GAG hinge between the coding sequence for NSP3 and the 2A element and a 3x FLAG (f) tag between the coding sequences for the 2A element and the S region. This approach was used to generate a set of vectors (collectively referred to as pT7/NSP3-CoV2/S vectors) that contained coding sequences for SARS-CoV-2 S1 (pT7/NSP3-2A-fS1), NTD (pT7/NSP3-2A-fNTD), RBD (pT7/NSP3-2A-fRBD), an extended form of the RBD (ExRBD) (pT7/NSP3-2A-fExRBD), and the S2 core region (CR) including its fusion domains (pT7/NSP3-2A-fCR) (Fig. 1). The S sequences were inserted into the pT7/NSP3SA11 vector at the same site as used before in the production of recombinant SA11 (rSA11) rotaviruses expressing FPs (15–17).

**Figure 2.**
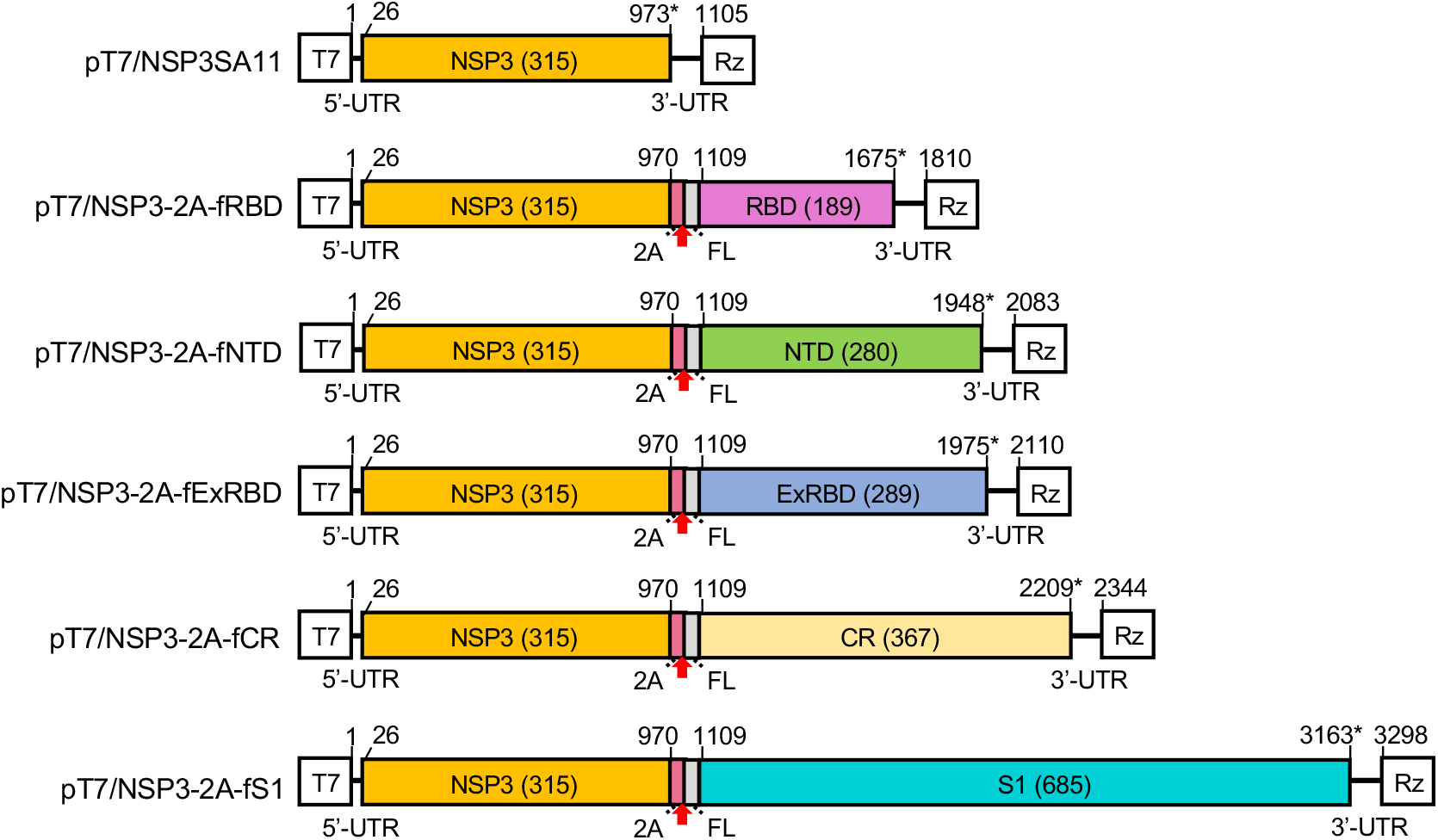
Plasmids with modified segment 7 (NSP3) cDNAs used to generate rSA11 viruses expressing regions of the SARS-CoV-2 S protein. Illustration indicates nucleotide positions of the coding sequences for NSP3, porcine teschovirus 2A element, 3xFLAG (FL), and the complete S1 or portions of the S1 (NTD, ExRBD, and RBD) and S2 (CR) proteins. The red arrow notes the position of the 2A translational stop-restart site, and the asterisk notes the end of the ORF. Sizes (aa) of encoded NSP3 and S products are in parenthesis. T7 (T7 RNA polymerase promoter sequence), Rz (Hepatitis D virus ribozyme), UTR (untranslated region).

### Recovery of rSA11 rotaviruses with segment 7 dsRNA containing S sequences

To generate rSA11 viruses, BHK-T7 monolayers were transfected with a complete set of pT7/SA11 expression vectors, except pT7/NSP3SA11 was replaced with a pT7/NSP3-CoV2/S vector, and a CMV expression plasmid (pCMV-NP868R) encoding the capping enzyme of African swine fever virus. In transfection mixtures, plasmids encoding rotavirus NSP2 (pT7/NSP2SA11) and NSP5 (pT7/NSP5SA11) were included at levels three-fold greater than the other pT7/SA11 vectors. BHK-T7 cells were overseeded with MA104 cells two days following transfection. The BHK-T7/MA104 cell mixture was freeze-thawed three days later, and the rSA11 viruses were recovered by plaque isolation and amplified by 1 or 2 cycles of growth in MA104 cells prior to characterization (36). Properties of the rSA11 viruses are summarized in Table 1.

**Table 1.**
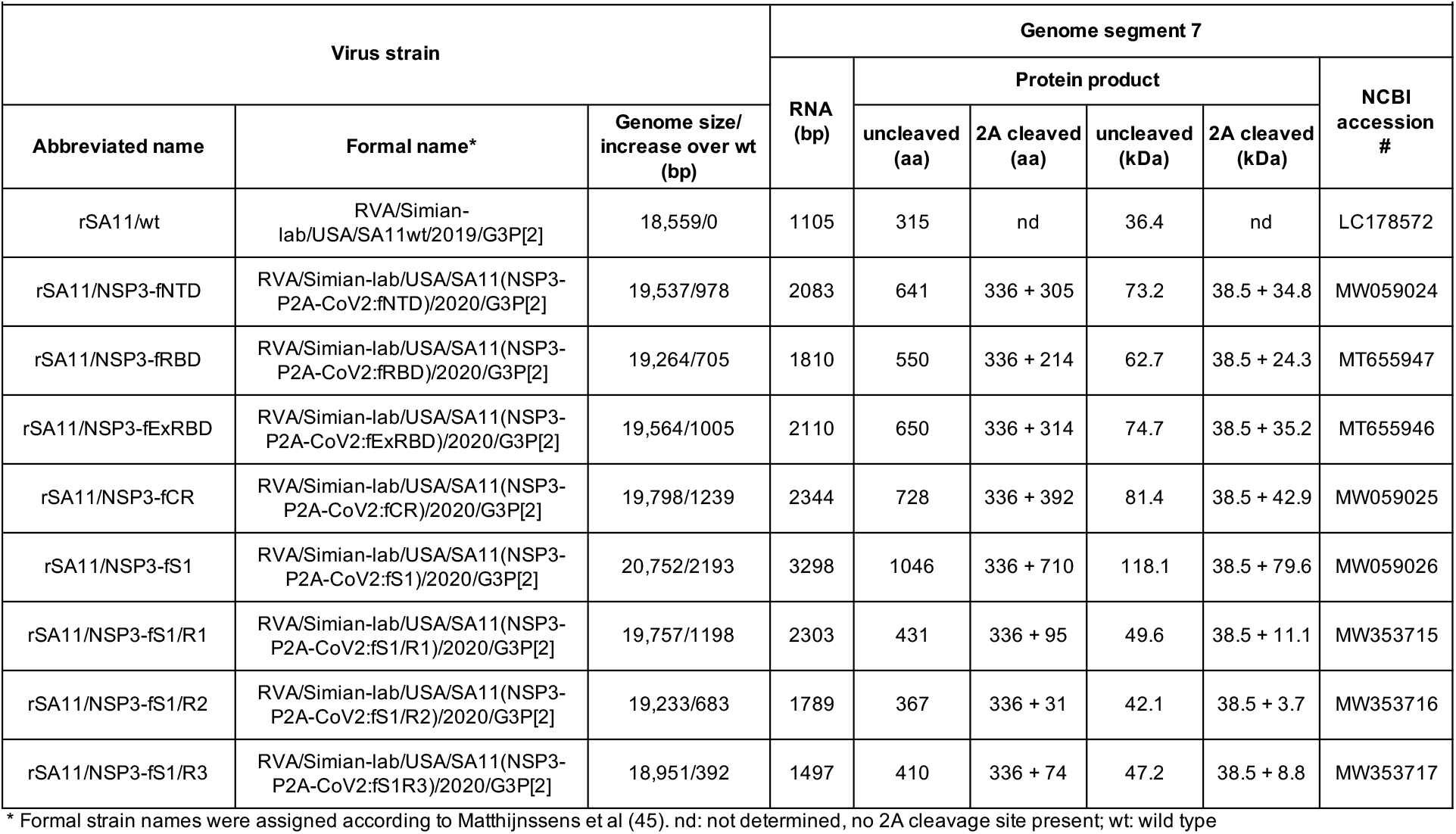
Properties of recombinant rRV/NSP3-2A-CoV2 strains.

Based on gel electrophoresis, rSA11 viruses generated with pT7/NSP3-S vectors (collectively referred to as rSA11/NSP3-CoV2/S viruses) contained segment 7 dsRNAs that were much larger than that of wildtype rSA11 (rSA11/wt) virus (Fig. 3). Sequence analysis confirmed that the segment 7 dsRNAs of the rSA11/NSP3-CoV2/S viruses matched the segment 7 sequences present in the pT7/NSP3-CoV2/S vectors (data not shown). The re-engineered segment 7 dsRNA of virus isolate rSA11/NSP3-fS1 had a length of 3.3 kbp, accounting for its electrophoretic migration near the largest rotavirus genome segment (segment 1), which is likewise 3.3 kbp in length (Table 1, Fig. 3A). The segment 7 dsRNA of rSA11/NSP3-fS1 contains a 2.2-kbp foreign sequence insertion, the longest foreign sequence that has been introduced into the segment 7 dsRNA, or for that matter, any rotavirus genome segment. The previously longest 7 dsRNA engineered into rSA11 was the 2.4-kbp segment 7 dsRNA of rSA11/NSP3-fmRuby-P2A-fUnaG, which contained a cassette that encoded three proteins (NSP3, UnaG, mRuby) (17). The total genome size of rSA11/NSP3-fS1 is 20.8 kbp, 12% greater than that of rSA11/wt (37). This is the largest genome known to exist within a rotavirus isolate and demonstrates the capacity of rotavirus to replicate and package large amounts of foreign sequence.

**Figure 3.**
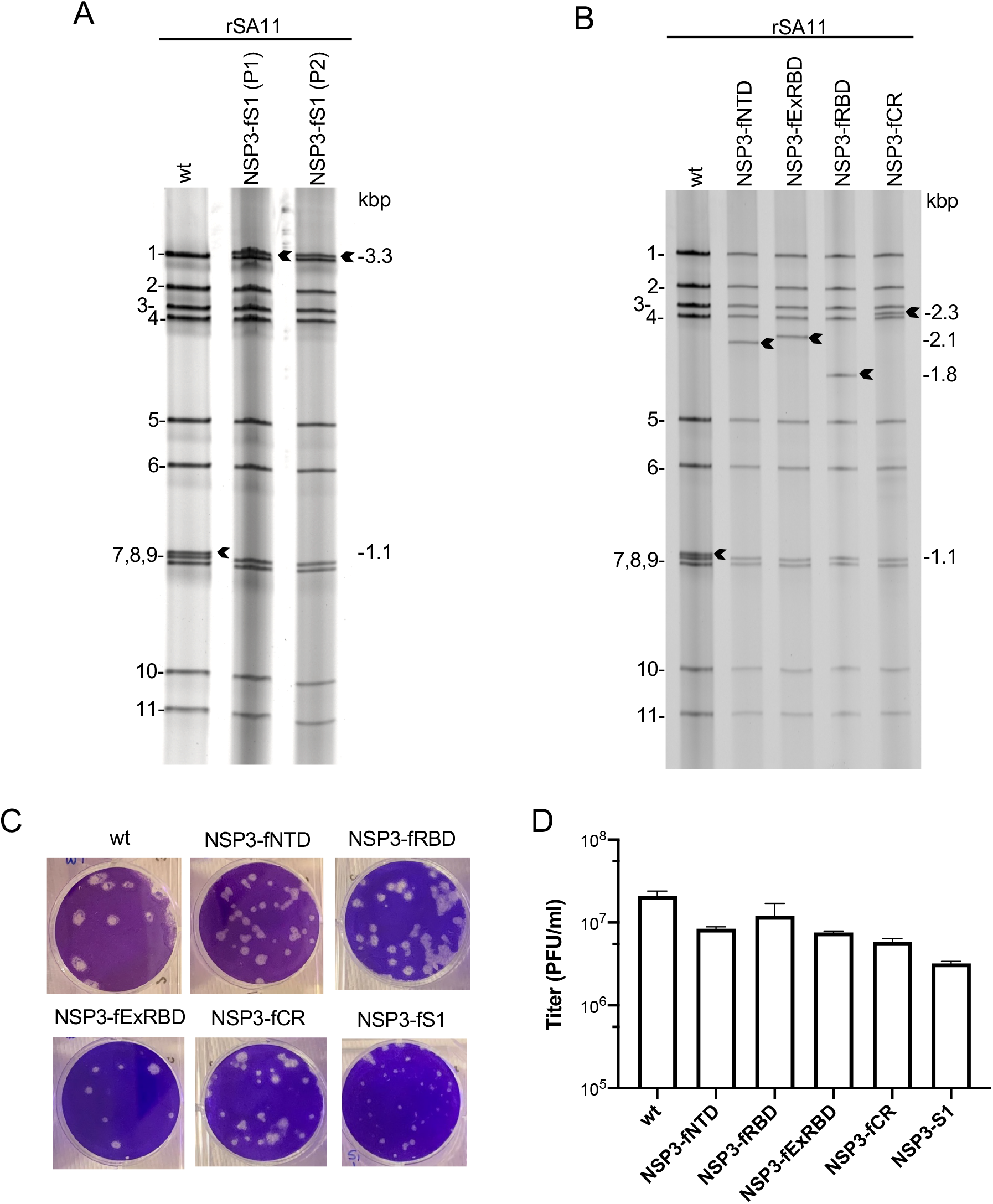
Properties of rSA11/NSP3-CoV2/S viruses expressing regions of the SARS-CoV-2 S protein. **(A and B)** dsRNA was recovered from MA104 cells infected with plaque-purified rSA11 isolates, resolved by gel electrophoresis, and detected by ethidium-bromide staining. RNA segments of rSA11/wt are labeled 1 to 11. Sizes (kbp) of segment 7 RNAs (black arrows) of rSA11 isolates are indicated. Double-stranded RNA of rSA11/NSP3-fS1 serially passaged twice (P1 and P2) in MA104 cells is shown in **(A). (C)** Plaque assays were performed using MA104 cells and detected by crystal-violet staining. **(D)** Titers reached by rSA11 isolates were determined by plaque assay. Bars indicate standard deviations calculated from three separate determinations.

The segment 7 dsRNAs of virus isolates, rSA11/NSP3-fNTD, -fRBD, -fExRBD, and -fCR, were determined to have lengths of 2.1, 1.8, 2.1, and 2.3 kbp, respectively (Table 1), and as expected from their sizes, migrated on RNA gels between rotavirus genome segments 3 (2.6 kbp) and 5 (1.6 kbp) (Fig. 3). The segment 7 dsRNAs of the rSA11/NSP3-fNTD, -fRBD, -fExRBD, and -fCR isolates contained foreign sequence insertions of 1.0, 0.7, 1.0, and 1.2 kbp, respectively, significantly smaller that the 2.1 kBP foreign sequence insertion of rSA11/NSP3-fS1. The smaller sizes of the foreign-sequence insert in the segment 7 RNAs of rSA11/NSP3-fNTD, -fRBD, -fExRBD, and -fCR provide additional genetic space that can be used to add routing and localization signals to S protein products, which may enhance their antigen processing and presentation, recognition by T cells, and trafficking to immune cells. For example, the extra genetic space can be used to add an N-terminal ER trafficking signal and a C-terminal plasma-membrane localization signal to the ExRBD, along with internal coiled-coil cassettes, that may favor surface presentation of a multimerized form of the ExRBD capable of inducing enhanced production of SARS-CoV-2 neutralizing antibodies.

Consistent with previous studies examining the phenotypes of rSA11 isolates expressing FPs (16–17), the sizes of plaques formed by rSA11/NSP3-CoV2/S viruses were smaller than plaques formed by rSA11/wt. Similarly, rSA11 viruses containing S-protein coding sequences grew to maximum titers that were up to 0.5-1 log lower than rSA11/wt. The reason for the smaller plaques and lower titers of the rSA11/NSP3-CoV2/S viruses is unknown, but may reflect the longer elongation time likely required for the viral RNA polymerase to transcribe their segment 7 dsRNAs during viral replication. Alternatively, it may reflect the longer time required to translate segment 7 (+)RNAs that contain S-protein coding sequences.

### Expression of S coding sequences by rSA11 rotaviruses

To determine whether the rSA11/NSP3-CoV2/S viruses expressed products from their S sequences, lysates prepared from MA104 cells infected with these viruses were examined by immunoblot assay using FLAG- and RBD-specific antibodies (Fig. 4A, B). Immunoblots probed with FLAG antibody showed that rSA11/NSP3-fNTD, -fExRBD, -fRBD, and -fCR viruses generated S products and that their sizes were as predicted for an active 2A element in the segment 7 ORF: fNTD (34.8 kDa), fExRBD (35.2 kDa), fRBD (24.3 kDa), and fCR (42.9 kDa) (Table 1). Immunoblot assays indicated that the rSA11/NSP3-fExRBD yielded higher levels of S product than any of the other rSA11/NSP3-CoV2/S viruses. The basis for the higher levels of the fExRBD product is unclear, but does not correlate with increased levels of expression of other viral products, such as NSP3 and VP6. Nonetheless, the high levels of ExRBD expression by rSA11/NSP3-fExRBD suggests that such viruses may be best suited in pursing the development of combined rotavirus/COVID vaccines.

**Figure 4.**
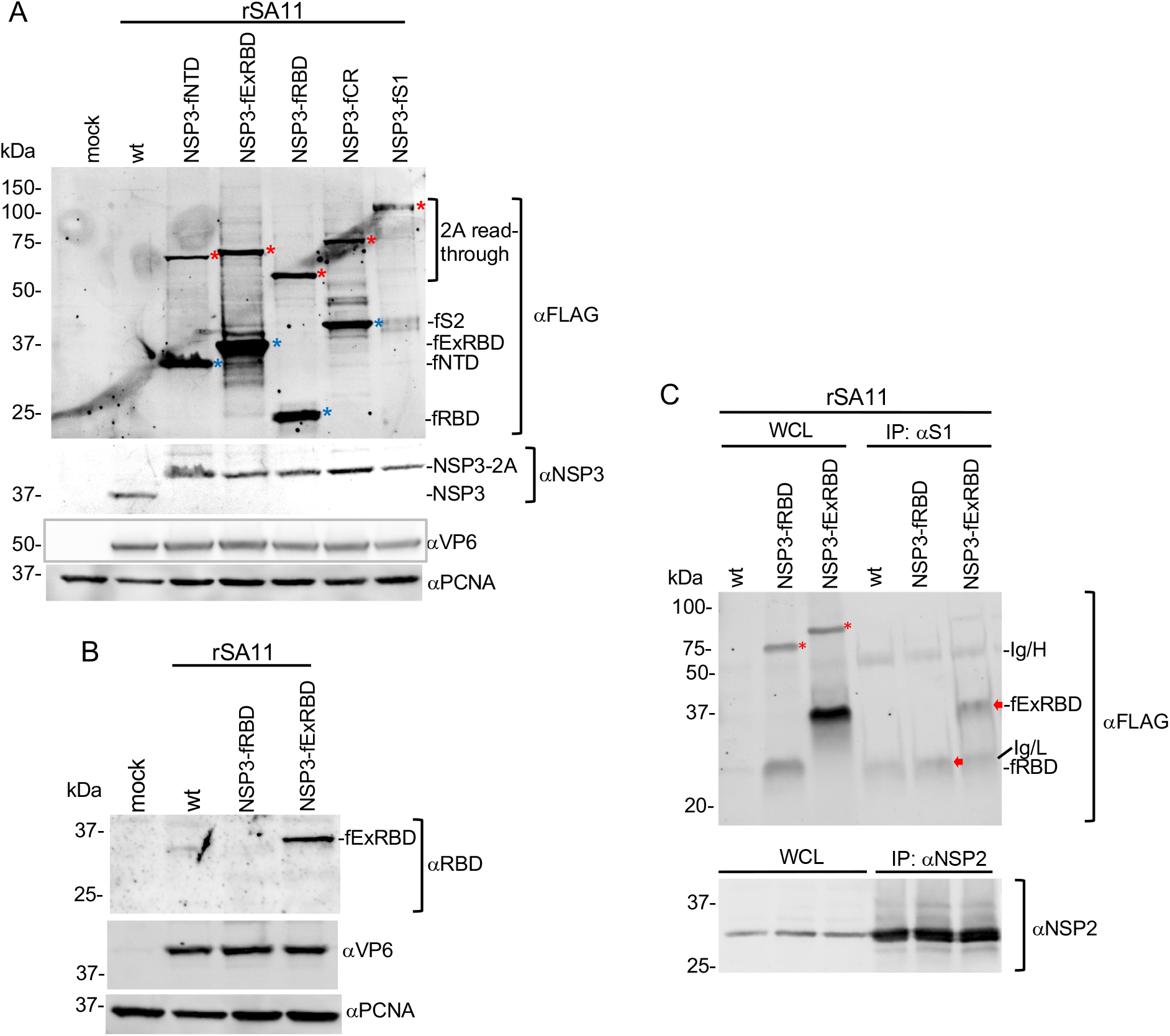
Expression of SARS-CoV-2 S products by rSA11 viruses. **(A, B)** Whole cell lysates (WCL) were prepared from cells infected with rSA11 viruses and examined by immunoblot assay using **(A)** FLAG antibody to detect S products (NTD, ExRBD, RBD, CR, S1, and 2A read-through products) and antibodies specific for rotavirus NSP3 and VP6 and cellular PCNA. Red asterisks identify 2A read-through products and blue asterisks identify 2A cleavage products. **(B)** Lysates prepared from MA104 cells infected with rSA11wt, rSA11/NSP3-fRBD and rSA11/NSP3-fExRBD were examined by immunoblot assay using antibodies specific for RBD (ProSci 9087), rotavirus VP6, and PCNA. **(C)** Lysates prepared from MA104 cells infected with rSA11/wt, rSA11/NSP3-fRBD and rSA11/NSP3-fExRBD viruses were examined by immunoprecipitation assay using a SARS-CoV-2 S1 specific monoclonal antibody (GeneTex CR3022). Lysates were also analyzed with a NSP2-specific polyclonal antibody. Antigen-antibody complexes were recovered using IgA/G beads, resolved by gel electrophoresis, blotted onto nitrocellulose membranes, and probed with FLAG (fRBD and fExRBD) and NSP2 antibody. Molecular weight markers are indicated (kDa). Red arrows indicate fRBD and fExRBD. fRBD comigrates near the Ig light chain (Ig/L). Ig heavy chain, Ig/H).

FLAG antibody did not detect the expected 79.6-kDa fS1 product in cells infected with rSA11/NSP3-fS1 (Fig. 4A). The S1 coding sequence in the segment 7 ORF includes an N-terminal signal sequence which, in SARS-CoV-2 infected cells, is cleaved from the S1 protein during synthesis on the endoplasmic reticulum (ER) (29,38). Cleavage of the signal sequence may have removed the upstream 3x FLAG tag from a S1 product, preventing its detection by the FLAG antibody. It is also possible that glycosylation and/or degradation of the 79.6 kDa-S1 product by ER-associated proteases may have prevented the protein’s detection. In addition, rotavirus which usurps and possibly remodels the ER in support of glycoprotein (NSP4 and VP7) synthesis and virus morphogenesis may perturb ER-interaction with the S signal sequence in such a way to prevent S1 synthesis (21). Interestingly, all the rSA11/NSP3-CoV2/S viruses, including rSA11/NSP3-fS1, generated 2A read-through products that were detectable using FLAG antibody. Thus, the 2A stop-start element in the rSA11/NSP3-2A-CoV2/S viruses was not fully active, which is consistent with previous reports analyzing the functionality of 2A elements within cells (39–41). However, with the exception of the rSA11/NSP3-fS1, all the viruses generated more 2A-cleaved S product than read-through product. Mutation of residues in and around the 2A element, including the inclusion of flexible linker sequences, may decrease the relative frequency of read through (42–43).

Lysates from MA104 cells infected with rSA11/wt, rSA11/NSP3-fRBD, and rSA11/NSP3-fExRBD were also probed with a RBD-specific polyclonal antibody prepared against a peptide mapping to the C-terminal end of the RBD domain (ProSci 9087). The RBD antibody recognized the fExRBD product of the rSA11/NSP3-fExRBD virus, but not the fRBD product of rSA11/NSP3-fRBD (Fig. 4B), presumably because the latter product lacked the peptide sequence used in generating the ProSci RBD antibody. To gain insight into whether the fRBD and fExRBD products folded into native structures mimicking those present in the SARS-CoV-2 S protein, lysates prepared from MA104 cells infected with rSA11/NSP3-fRBD and rSA11/NSP3-fExRBD were probed by pulldown assay using an anti-RBD conformation-dependent neutralizing monoclonal antibody (GeneTex CR3022). As shown in Fig. 4C, the CR3022 immunoprecipitate included fExRBD, indicating that this product included a neutralizing epitope found in authentic SARS-CoV-2 S protein. Thus, at least some of the RBD product of rSA11/NSP3-fExRBD has likely folded in a conformation capable of inducing a protective antibody response. Unlike the successful pulldown of ExRBD with CR3022 antibody, it was not clear if the antibody likewise immunoprecipitated the fRBD product of rSA11/NSP3-fRBD. This uncertainty stems from the light chain of the CR3022 antibody obscuring the electrophoretically closely-migrating fRBD product in immunoblot assays (Fig. 4C).

### Expression of the ExRBD and RBD products by rSA11s during rotavirus infection

To gain insight into fExRBD and fRBD expression during virus replication, MA104 cells were infected with rSA11/wt, rSA11/NSP3-fExRBD or rSA11/NSP3-fRBD and then harvested at intervals between 0 and 12 hr p.i. Analysis of the infected cell lysates by immunoblot assay showed that fExRBD and fRBD were readily detectable by 4 h p.i., paralleling the expression of rotavirus proteins NSP3 and VP6 (Fig. 5). Increased levels of fExRBD and fRBD were present at 8 and 12 h p.i., without obvious accumulation of FLAG-tagged products of smaller sizes. Thus, the fExRBD and fRBD products appear to be relatively stable.

**Figure 5.**
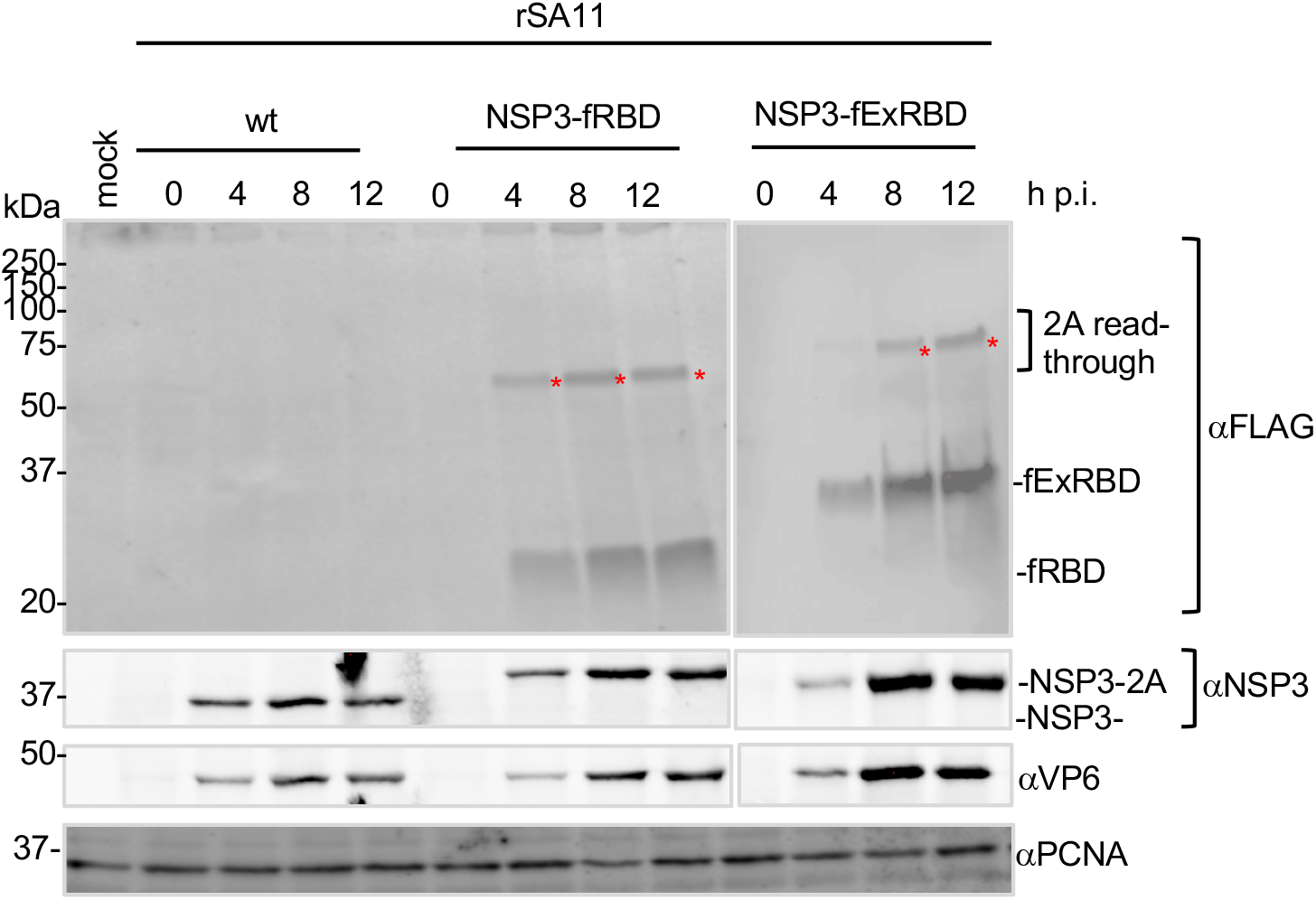
Production of RBD and ExRBD by rSA11 viruses during infection. MA104 cells were mock infected or infected with rSA11/wt, rSA11/NSP3-fRBD, or rSA11/NSP3-fExRBD (MOI of 5). Lysates were prepared from the cells at 0, 4, 8, or 12 h p.i. and analyzed by immunoblot assay using antibodies specific for FLAG, NSP3, VP6, and PCNA. Red asterisks identify 2A read-through products. Positions of molecular weight markers are indicated (kDa).

### Density of rSA11 virus particles containing S sequences

The introduction of S sequences into the rSA11/NSP3-CoV2/S viruses increased the size of their viral genomes by 1.0 to 2.5 kbp beyond that of SA11/wt. Assuming the rSA11/NSP3-CoV2/S viruses are packaged efficiently and contain a complete constellation of 11 genome segments, the increased content of dsRNA within the core of rSA11/NSP3-CoV2/S particles should cause their densities to be greater than that of SA11/wt particles. To explore this possibility, rSA11/wt (18.6-kbp genome), rSA11/NSP3-fExRBD (19.5 kbp) and rSA11/NSP3-fS1 (20.8 kbp) were amplified in MA104 cells. The infected-cell lysates were then treated with EDTA to convert rotavirus virions (triple-layered particles) into double-layered particles (DLPs). The particles were centrifuged to equilibrium on CsCl gradients (Fig. 6) and the density of the DLP bands determined by refractometry. The analysis indicated that the density of rSA11/NSP3-fExRBD DLPs (1.386 g/cm^3^) was greater than SA11/wt DLPs (1.381 g/cm^3^) (panel A) and similarly, the density of rSA11/NSP3-fS1 DLPs (1.387 g/cm^3^) was greater that SA11/wt DLPs (1.38 g/cm^3^) (panel B). Analysis of the banded DLPs by gel electrophoresis confirmed that they contained the expected constellation of eleven genome segments. To confirm that the density of rSA11/NSP3-fS1 DLPs was different that rSA11/wt DLPs, infected-cell lysates containing each of these viruses were pooled, treated with EDTA, and the viral DLPs in the combined sample banded by centrifugation on a CsCl gradient (Fig. 6, panel E). Analysis of the gradient revealed the presence of two bands of particles, indicating that rSA11/NSP3-fSA11-fS1 and rSA11/wt DLPs were of different densities. Gel electrophoresis of the combined DLP bands showed, as expected, that both rSA11/NSP3-fSA11-fS1 and rSA11/wt were present. Taken together, these results demonstrate that rSA11/NSP3-CoV-2/S virions contain complete genome constellations despite the fact that their genome sizes are significantly greater than that of wildtype SA11 virus. Indeed, the 20.8-kbp rSA11/NSP3-fS1 genome is 12% greater in size than the 18.6-kbp rSA11/wt genome (Table 1). Thus, the rotavirus core has space to accommodate large amounts of additional foreign sequence. How the dsRNA within the core is re-distributed to accommodate large amounts of additional sequence is not known, but clearly the core remains a transcriptionally-active nanomachine despite the additional sequence. Whether other genome segments can be engineered similarly to segment 7 of rSA11/NSP3-fS1 to include 2 kb of additional sequence remains to be determined. The maximum packaging capacity of the core also remains to be determined.

**Figure 6.**
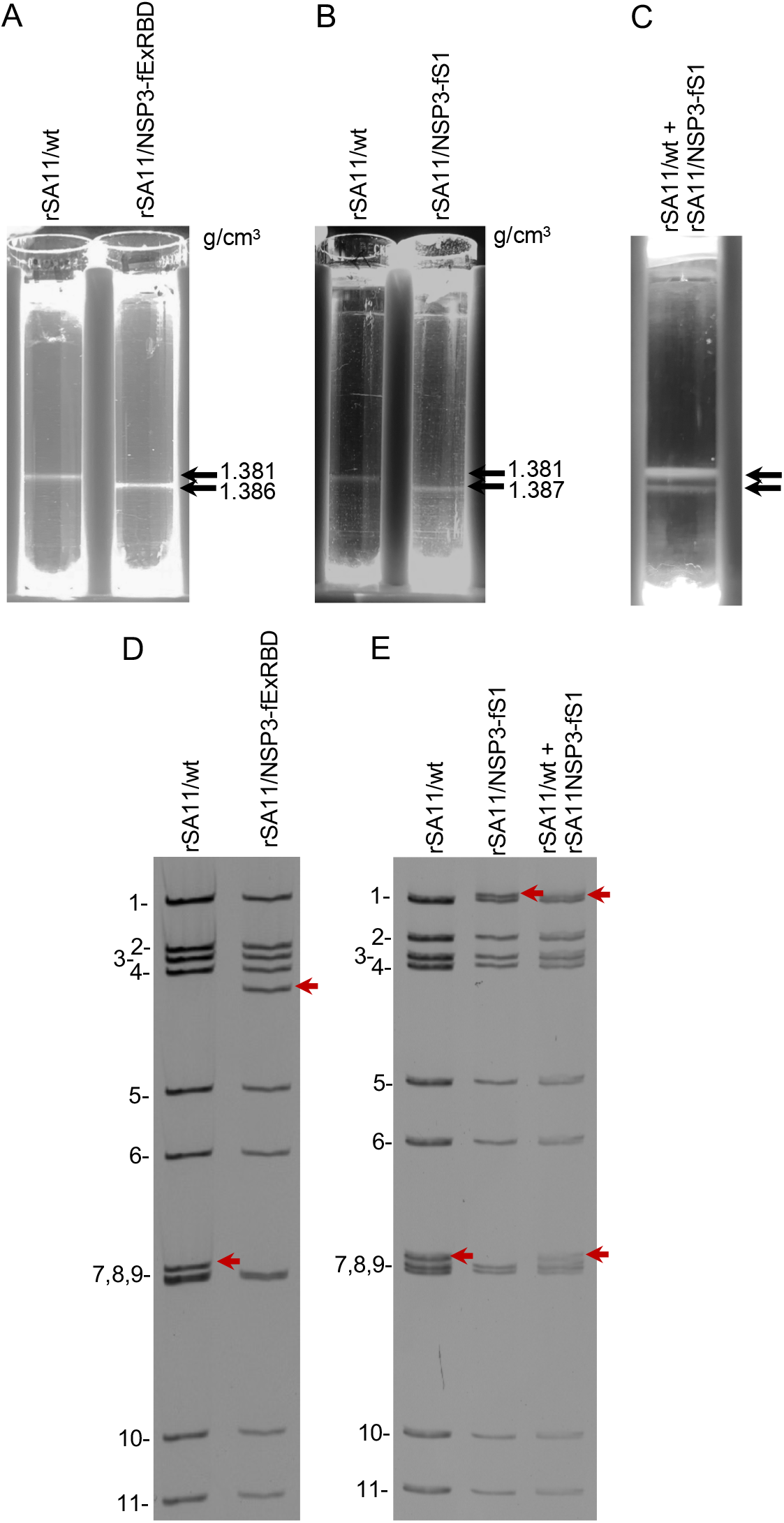
Impact of genome size on rotavirus particle density. MA104 cells were infected with rSA11/wt, rSA11/NSP3-fExRBD, or rSA11/NSP3-fS1 viruses at an MOI of 5. At 12 h p.i., the cells were recovered, lysed by treatment with non-ionic detergent, and treated with EDTA to convert rotavirus virions into DLPs. **(A, B)** DLPs were banded by centrifugation in CsCl gradients and densities (g/cm^3^) were determined using a refractometer. **(C)** Lysates from rSA11/wt and rSA11/NSP3-fS1 infected cells were combined and their DLP components banded by centrifugation in a CsCl gradient. **(D, E)** Electrophoretic profile of the dsRNA genomes of DLPs recovered from CsCl gradients. Panel D RNAs derive from DLPs in panel A and panel E RNAs derive from DLPs in panel B and C. RNA segments of rSA11/wt are labeled 1 to 11. Positions of segment 7 RNAs are indicated with red arrows.

**Figure 7.**
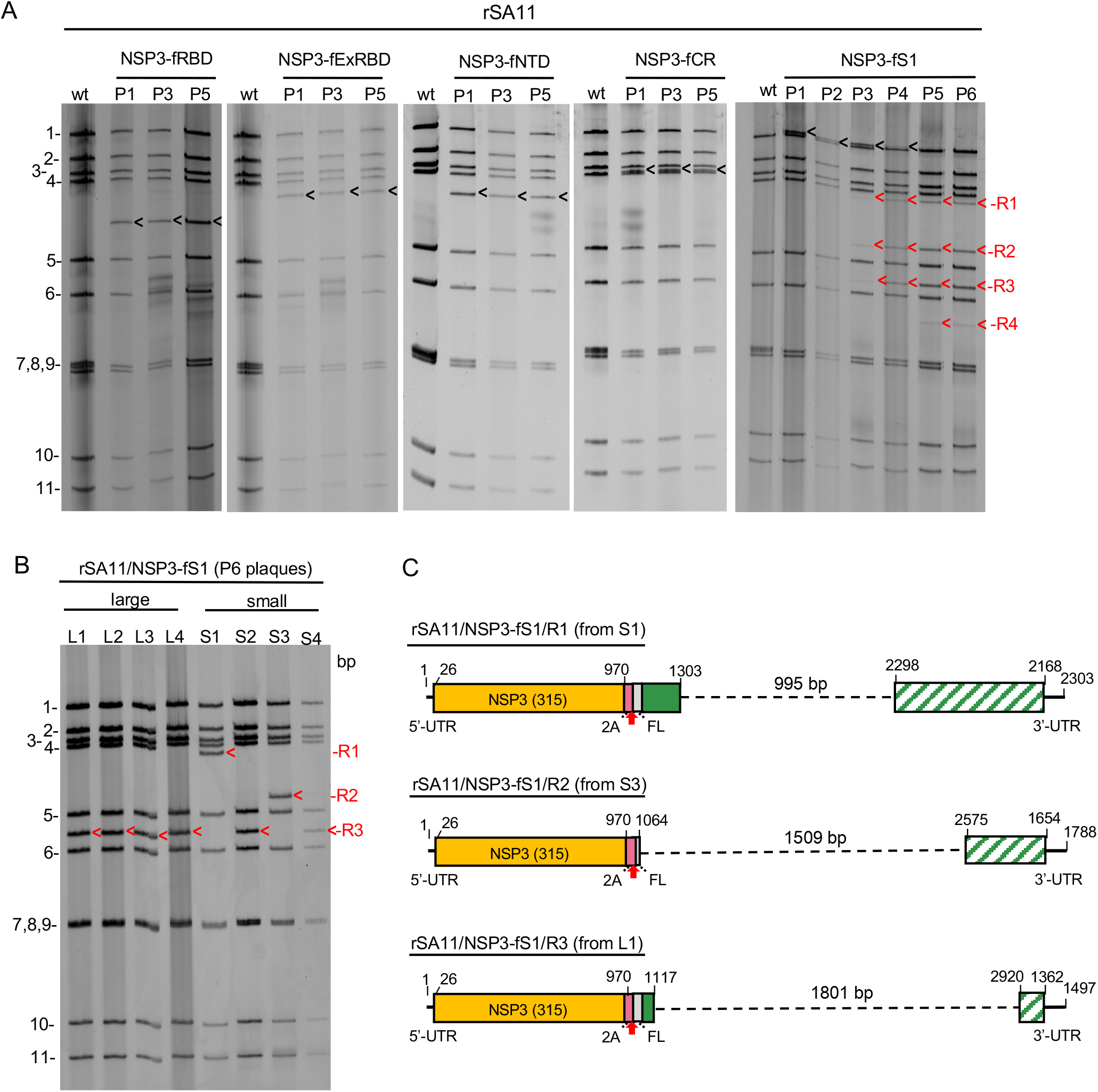
Genetic stability of rSA11 strains expressing SARS-CoV-2 S domains. rSA11 strains were serially passaged 5 to 6 times (P1 to P5 or P6) in MA104 cells. **(A)** Genomic RNAs were recovered from infected cell lysates and analyzed by gel electrophoresis. Positions of viral genome segments are labeled. Position of modified segment 7 (NSP3) dsRNAs introduced into rSA11 strains are denoted with black arrows. Genetic instability of the modified segment 7 (NSP3) dsRNA of rSA11/NSP3-fS1 yielded R1-R4 RNAs during serial passage. **(B)** Genomic RNAs prepared from large (L1-L4) and small (S1-S4) plaque isolates of P6 rSA11/NSP3-fS1. Segment 7 RNAs are identified as R1-R3, as in **(A)**. **(C)** Organization of R1-R3 sequences determined by sequencing of segment 7 RNAs of L1, S1, and S3 plaque isolates. Sequence deletions are indicated with dashed lines. Regions of the S1 ORF that are no longer encoded by the R1-R3 segment 7 RNAs are indicated by slashed green-white boxes.

### Genetic stability of rSA11 rotaviruses containing S sequences

The genetic stability of the rSA11/NSP3-CoV2/S viruses were assessed by serial passage, with a fresh monolayer of MA104 cells infected with 1:1000 dilutions of cell lysates at each round. Electrophoretic analysis of the dsRNAs recovered from cells infected with rSA11/NSP3-fNTD, -fRBD, -ExRBD, or -ExCR showed no changes in the sizes of any of the 11 genome segments over 5 rounds of passage (P1-P5), including segment 7, indicating that these viruses were genetically stable. In contrast, serial passage of rSA11/NSP3-S1 showed evidence of instability. By the third round of passage, novel genome segments were appearing that were smaller than the 3.3-kbp segment 7 RNA. With continued passage, four novel segments (R-1 to R-4) became prominent and the 3.3-kbp segment 7 RNA was no longer detectable, suggesting that the high-passage virus pools (P3-P6) were populated by variants containing segment 7 RNAs derived from the 3.3-kb segment 7 RNA through internal sequence deletion. To evaluate this possibility, 8 variants were recovered from the P6 virus pool by plaque isolation, 4 with a large (L) plaque phenotype and 4 with a small (S) plaque phenotype. Electrophoretic analysis of the genomes of the variants showed that none contained the 3.3-kbp segment 7 RNA. Instead, 6 variants (L1, L2, L3, L4, S2, and S4) contained the R3 segment, and the other two variants contained either the R1 (S1) or R2 (R2) segment. No variants were recovered that contained the novel R4 segment. Sequencing showed that the R1, R2, and R3 segments were in fact derivatives of the 3.3-kbp segment 7 RNA. The R1, R2, and R3 RNAs all retained the complete 5’- and 3’-UTRs and NSP3 ORF of segment 7, but contained sequence deletions of 1.0 (R1), 1.5 (R2), or 1.8 (R3) kbp of S1 coding sequence. The fact that 6 of the 8 variants isolated by plaque assay contained the R3 segment suggests that variants with this RNA may have a growth advantage over variants with the R1, R2, or R4 RNAs. Although genetic instability gave rise to rSA11/NSP3-fS1 variants lacking portions of the S1 ORF, none were identified that lacked portions of the NSP3 ORF. This suggests that NSP3 may be essential for virus replication, which would explain the failure of previous efforts by us to recover viable rSA11s encoding truncated forms of NSP3 through insertion of stop codons in the NSP3 ORF (data not shown).

### Summary

We have shown that reverse genetics can be used to generate recombinant rotaviruses that express, as separate products, portions of the SARS-CoV-2 S protein, including its immunodominant RBD. These results indicate that it may be possible to develop rotaviruses as vaccine expression vectors, providing a path for generating oral live-attenuated rotavirus-COVID-19 combination vaccines able to induce immunological protective responses against both rotavirus and SARS-CoV-2. Such combination vaccines would be designed for use in infants and young children and would allow the widespread distribution and administration of COVID-19-targeted vaccines by piggy backing onto current rotavirus immunization programs used in the USA and many other countries, both developed and developing. In addition, our findings raise the possibility that through the use of rotavirus as vaccine expression platforms, rotavirus-based combination vaccines could be made against other enteric viruses including norovirus, astrovirus, and hepatitis E virus.

We have determined that the 18.6-kbp rotavirus dsRNA can accommodate as much as 2.2-kbp of foreign sequence, which is sufficient to encode the SARS-CoV-2 S1 protein. However, in our hands, rSA11s encoding S1 were not genetically stable and failed to express the appropriate S1 product, for reasons that are uncertain but under further investigation. Rotaviruses carrying large amounts of foreign sequence are characteristically genetically unstable (this study and data not shown), but those with foreign sequences of <1.0-1.5-kbp are stable over 5-10 rounds of serial passage at low MOI and, thus, can be developed into vaccine candidates. The coding capacity provided by 1.0-1.5-kbp of extra sequence is sufficient to produce recombinant rotaviruses that encode the SARS-CoV-2 NTD, RBD, or S2 core along with trafficking signals that can promote engagement of S products with antigen-presenting cells and naive B-lymphocytes. Current work is underway to gain insight how successful rotaviruses expressing SARS-CoV-2 products are in inducing neutralizing antibodies in immunized animals.

## MATERIALS AND METHODS

### Cell culture

Embryonic monkey kidney cells (MA104) were grown in medium 199 (M199) containing 5% fetal bovine serum (FBS) and 1% penicillin-streptomycin. Baby hamster kidney cells expressing T7 RNA polymerase (BHK-T7) were provided by Dr. Ulla Buchholz, Laboratory of Infectious Diseases, NIAID, NIH, and were propagated in Glasgow minimum essential media (GMEM) containing 5% heat-inactivated fetal bovine serum (FBS), 10% tryptone-peptide broth, 1% penicillin-streptomycin, 2% non-essential amino acids, and 1% glutamine (36). BHK-T7 cells were grown in medium supplemented with 2% Geneticin (Invitrogen) with every other passage.

### Plasmid construction

Recombinant SA11 rotaviruses were prepared using the plasmids pT7/VP1SA11, pT7/VP2SA11, pT7/VP3SA11, pT7/VP4SA11, pT7/VP6SA11, pT7/VP7SA11, pT7/NSP1SA11, pT7/NSP2SA11, pT7/NSP3SA11, pT7/NSP4SA11, and pT7/NSP5SA11 [https://www.addgene.org/Takeshi_Kobayashi/] and pCMV-NP868R (16). The plasmid pT7/NSP3-P2A-fUnaG was produced, as described elesewhere, by fusing a DNA fragment containing the ORF for P2A-3xFL-UnaG to the 3’-end of the NSP3 ORF in pT7/NSP3SA11 (17). A plasmid (pTWIST/COVID19spike) containing a full-length cDNA of the SARS-CoV-2 S gene (GenBank MN908947.3) was purchased from Twist Bioscience. The plasmids pT7/NSP3-2A-fNTD, pT7/NSP3-2A-fExRBD, pT7/NSP3-2A-fRBD, pT7/NSP3-2A-fCR, and pT7/NSP3-2A-S1 were made by replacing the UnaG ORF in pT7/NSP3-2A-fUnaG with ORFs for the NTD, ExRBD, RBD, CR, and S1 regions, respectively, of the SARS-CoV-2 S protein, by In-Fusion cloning. DNA fragments containing NTD, ExRBD, RBD, CR, and S1 coding sequences were amplified from pTWIST/COVID19spike using the primer pairs NTD_For and NTD_Rev, ExRBD_For and ExRBD_Rev, RBD_For and RBD_Rev, CR_For and CR_Rev, and S1_For and S1_Rev, respectively (Table 2). Transfection quality plasmids were prepared commercially (www.plasmid.com) or using Qiagen plasmid purification kits. Primers were provided by and sequences determined by EuroFins Scientific.

**Table 2.**
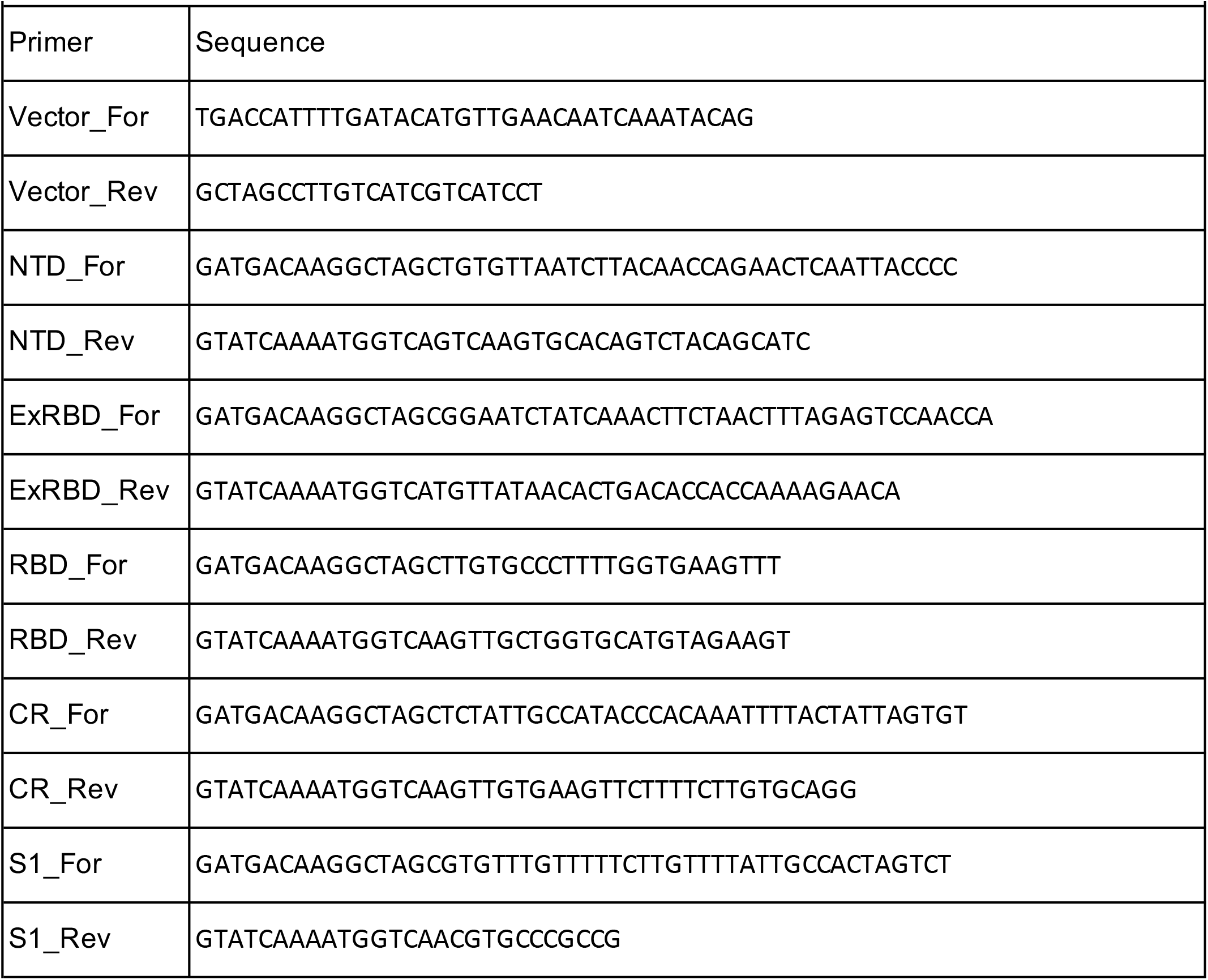
Primers used to produce pT7/NSP3-2A-CoV2 plasmids.

### Recombinant viruses

The reverse genetics protocol used to generate recombinant rotaviruses was described in detail previously (16,44). To summarize, BHK-T7 cells were transfected with SA11 pT7 plasmids and pCMV-NP868R using Mirus TransIT-LT1 transfection reagent. Two days later, the transfected cells were overseeded with MA104 cells and the growth medium (serum-free) adjusted to a final concentration of 0.5 μg/ml trypsin. Three days later, the BHK-T7/MA104 cell mixture was freeze-thawed 3-times and the lysates clarified by low-speed centrifugation. Recombinant virus in clarified lysates were amplified by one or two rounds of passage in MA104 cells maintained in serum-free medium containing 0.5 μg/ml trypsin. Individual virus isolates were obtained by plaque purification and typically amplified 1 or 2 rounds in MA104 cells prior to analysis. Viral dsRNAs were recovered from infected-cell lysates by Trizol extraction, resolved by electrophoresis on Novex 8% polyacrylamide gels (Invitrogen) in Tris-glycine buffer, and detected by staining with ethidium bromide. Viral dsRNAs in gels were visualized using a BioRad ChemiDoc MP Imaging System. The genetic stability of plaque isolated rSA11s was assessed by serial passage as described previously (17).

### Immunoblot analysis

MA104 cells were mock infected or infected with 5 PFU of recombinant virus per cell and harvested at 8 h p.i. Cells were washed with cold phosphate-buffered saline (PBS), pelleted by low-speed centrifugation, and lysed by resuspending in lysis buffer [300 mM NaCl, 100 mM Tris-HCl, pH 7.4, 2% Triton X-100, and 1x EDTA-free protease inhibitor cocktail (Roche cOmplete)]. For immunoblot assays, lysates were resolved by electrophoresis on Novex linear 8-16% polyacrylamide gels and transferred to nitrocellulose membranes. After blocking with phosphate-buffered saline containing 5% non-fat dry milk, blots were probed with guinea pig polyclonal NSP3 (Lot 55068, 1:2000) or VP6 (Lot 53963, 1:2000) antisera (2), mouse monoclonal FLAG M2 (Sigma F1804, 1:2000), rabbit monoclonal PCNA [13110S, Cell Signaling Technology (CST), 1:1000] antibody or rabbit anti-RBD (ProSci 9087; 1:200) antibody. Primary antibodies were detected using 1:10,000 dilutions of horseradish peroxidase (HRP)-conjugated secondary antibodies: horse anti-mouse IgG (CST), anti-guinea pig IgG (KPL), or goat anti-rabbit IgG (CST). Signals were developed using Clarity Western ECL Substrate (Bio-Rad) and detected using a Bio-Rad ChemiDoc imaging system.

### Immunoprecipitation assay

Mock-infected and infected cell lysates were prepared as above. Lysates were mixed with a SARS-CoV-2 S1 specific monoclonal antibody (GeneTex CR3022, 1:150 dilution) or an NSP2 monoclonal antibody (#171, 1:200). After incubation at 4C with gentle rocking for 18 h, antigen-antibody complexes were recovered using Pierce magnetic IgA/IgG beads (ThermoFisher Scientific), resolved by gel electrophoresis, and blotted onto nitrocellulose membranes. Blots were probed with FLAG antibody (1:2000) to detect fRBD and fExRBD and NSP2 antibody (1:2000).

### CsCl gradient centrifugation

MA104-cell monolayers in 10-cm cell culture plates were infected with rSA11 viruses at an MOI of 5 and harvested at 12 h p.i. Cells were lysed by adjusting media to 0.5% Triton X100 (Sigma) and incubation on ice for 5 min. Lysates were then clarified by centrifugation at 500 x g at 4C for 6 min. The clarified lysates were adjusted to 10 mM EDTA and incubated for 1 h at 37C to cause the conversion of rotavirus TLPs to DLPs (36). CsCl was added to samples to a density of 1.367 g/cm^3^ and samples were centrifuged at 110,000 x *g* with a Beckman SW55Ti rotor at 8C for 22 h. Fractions containing viral bands were recovered using a micropipettor and fraction densities were determined using a refractometer.

### Genetic stability of rSA11 viruses

Viruses were serially passaged on MA104-cell monolayers using 1:1000 dilutions of infected cell lysates prepared in serum-free M199 medium and 0.5 μg/ml trypsin. When cytopathic effects reached completion (4-5 days), cells were freeze-thawed twice in their medium, and lysates were clarified by low-speed centrifugation. To recover dsRNA, clarified lysates (600 ul) were extracted with Trizol (ThermoFisher Scientific). The RNA samples were resolved by electrophoresis on 8% polyacrylamide gels and the bands of dsRNA detected by ethidium-bromide staining.

### GenBank accession numbers

Segment 7 sequences in rSA11 viruses have been deposited in Genbank: wt (LC178572), NSP3-P2A-fNTD (MW059024), NSP3-P2A-fRBD (MT655947), NSP3-P2A-ExRBD (MT655946), NSP3-P2A-fCR (MW059025), NSP3-P2A-S1 (MW059026), NSP3-P2A-S1/R1 (MW353715), NSP3-P2A-S1/R2 (MW353716), and NSP3-P2A-S1/R3 (MW353717). See also Table 1.

## ACKNOWLEDGEMENT

Our thanks go to lab members for their support and encouragement on this project. This work was funded by National Institutes of Health grant R21AI144881, Indiana University Start-Up Funding, and the Lawrence M. Blatt Endowment.

## References

1. Dai L, Gao GF. 2021. Viral targets for vaccines against COVID-19. Nat Rev Immunol 21, 73–82.

2. Kaur SP, Gupta V. 2020. COVID-19 Vaccine: A comprehensive status report. Virus Research 288: 198114.

3. Ludvigsson JF. 2020. Systematic review of COVID-19 in children shows milder cases and a better prognosis than adults. Acta Paediatr 109(6):1088–1095.

4. Pollán M, Pérez-Gómez B, Pastor-Barriuso R, Oteo J, Hernán MA, Pérez-Olmeda M, Sanmartín JL, Fernández-García A, Cruz I, Fernández de Larrea N, Molina M, Rodríguez-Cabrera F, Martín M, Merino-Amador P, León Paniagua J, Muñoz-Montalvo JF, Blanco F, Yotti R; ENE-COVID Study Group. 2020. Prevalence of SARS-CoV-2 in Spain (ENE-COVID): a nationwide, population-based seroepidemiological study. Lancet 396 (10250): 535–544.

5. Burke RM, Tate JE, Kirkwood CD, Steele AD, Parashar UD. 2019. Current and new rotavirus vaccines. Curr Opin Infect Dis 32: 435–444.

6. Folorunso OS, Sebolai OM. 2020. Overview of the development, impacts, and challenges of live-attenuated oral rotavirus vaccines. Vaccines (Basel) 8 (3): 341.

7. Ali A, Kazi AM, Cortese MM, Fleming JA, Moon S, Parashar UD, Jiang B, McNeal MM, Steele D, Bhutta Z, Zaidi AK. 2015. Impact of withholding breastfeeding at the time of vaccination on the immunogenicity of oral rotavirus vaccine--a randomized trial. PLoS One 10 (6): e0127622. Erratum in: PLoS One 10 (12): e0145568.

8. Liu GF, Hille D, Kaplan SS, Goveia MG. 2017. Postdose 3 G1 serum neutralizing antibody as correlate of protection for pentavalent rotavirus vaccine. Hum Vaccin Immunother 13 (10): 2357–2363.

9. Angel J, Steele AD, Franco MA. 2014. Correlates of protection for rotavirus vaccines: Possible alternative trial endpoints, opportunities, and challenges. Hum Vaccin Immunother 10 (12): 3659–71.

10. Leshem E, Tate JE, Steiner CA, Curns AT, Lopman BA, Parashar UD. 2015. Acute gastroenteritis hospitalizations among US children following implementation of the rotavirus vaccine. JAMA. 2015 313 (22): 2282–4. Erratum in: JAMA 314 (2): 188.

11. Clark A, Black R, Tate J, Roose A, Kotloff K, Lam D, Blackwelder W, Parashar U, Lanata C, Kang G, Troeger C, Platts-Mills J, Mokdad A; Global Rotavirus Surveillance Network, Sanderson C, Lamberti L, Levine M, Santosham M, Steele D. 2017. Estimating global, regional and national rotavirus deaths in children aged <5 years: Current approaches, new analyses and proposed improvements. PLoS One 12 (9): e0183392.

12. Kanai Y, Kawagishi T, Nouda R, Onishi M, Pannacha P, Nurdin JA, Nomura K, Matsuura, Y, Kobayashi T. 2018. Development of stable rotavirus reporter expression systems. J Virol 93: e01774–18.

13. Komoto S, Fukuda S, Ide T, Ito N, Sugiyama M, Yoshikawa T, Murata T, Taniguchi K. 2018. Generation of recombinant rotaviruses expressing fluorescent proteins by using an optimized reverse genetics system. J Virol 92: e00588–18.

14. Sánchez-Tacuba L, Feng N, Meade NJ, Mellits KH, Jaïs PH, Yasukawa LL, Resch TK, Jiang B, López S, Ding S, Greenberg HB. 2020. An optimized reverse genetics system suitable for efficient recovery of simian, human, and murine-like rotaviruses. J Virol 94 (18): e01294–20.

15. Philip AA, Herrin BE, Garcia ML, Abad AT, Katen SP, Patton JT. 2019. Collection of recombinant rotaviruses expressing fluorescent reporter proteins. Microbio Resour Announc 8 (27): e00523–19.

16. Philip AA, Perry JL, Eaton HE, Shmulevitz M, Hyser JM, Patton JT. 2019. Generation of recombinant rotavirus expressing NSP3-UnaG fusion protein by a simplified reverse genetics system. J Virol 93: e01616–19.

17. Philip AA, Patton JT. 2020. Expression of separate heterologous proteins from the rotavirus NSP3 genome segment using a translational 2A stop-restart element. J Virol 94 (18): e00959–20.

18. Kanai Y, Komoto S, Kawagishi T, Nouda R, Nagasawa N, Onishi M, Matsuura Y, Taniguchi K, Kobayashi T. 2017. Entirely plasmid-based reverse genetics system for rotaviruses. Proc Natl Acad Sci USA 114: 2349–2354.

19. Komoto S, Fukuda S, Kugita M, Hatazawa R, Koyama C, Katayama K, Murata T, Taniguchi K. 2019. Generation of infectious recombinant human rotaviruses from just 11 cloned cDNAs encoding the rotavirus genome. J Virol 93 (8): e02207–18.

20. Crawford SE, Ramani S, Tate JE, Parashar UD, Svensson L, Hagbom M, Franco MA, Greenberg HB, O’Ryan M, Kang G, Desselberger U, Estes MK. 2017. Rotavirus infection. Nat Rev Dis Primers 3: 17083.

21. Trask SD, McDonald SM, Patton JT. 2012. Structural insights into the coupling of virion assembly and rotavirus replication. Nat Rev Microbiol 10: 165–177.

22. Eaton HE, Kobayashi T, Dermody TS, Johnston RN, Jais PH, Shmulevitz M 2017. African swine fever virus NP868R capping enzyme promotes reovirus rescue during reverse genetics by promoting reovirus protein expression, virion assembly, and RNA incorporation into infectious virions. J Virol 91: e02416–16.

23. Criglar JM, Crawford SE, Zhao B, Smith HG, Stossi F, Estes MK. 2020. A genetically engineered rotavirus NSP2 phosphorylation mutant impaired in viroplasm formation and replication shows an early interaction between vNSP2 and cellular lipid droplets. J Virol 94 (15): e00972–20.

24. Navarro, A., Trask, S.D., Patton, J.T. 2013. Generation of genetically stable recombinant rotaviruses containing novel genome rearrangements and heterologous sequences by reverse genetics. J Virol 87: 6211–6220.

25. Chang-Graham AL, Perry JL, Strtak AC, Ramachandran NK, Criglar JM, Philip AA, Patton JT, Estes MK, Hyser JM. 2019. Rotavirus calcium dysregulation manifests as dynamic calcium signaling in the cytoplasm and endoplasmic reticulum. Sci Rep 9 (1):10822.

26. Komoto S, Kanai Y, Fukuda S, Kugita M, Kawagishi T, Ito N, Sugiyama M, Matsuura Y, Kobayashi T, Taniguchi K. 2017. Reverse genetics system demonstrates that rotavirus nonstructural protein NSP6 is not essential for viral replication in cell culture. J Virol 91: e00695–17.

27. Gratia M, Sarot E, Vende P, Charpilienne A, Baron CH, Duarte M, Pyronnet S, Poncet D. 2015. Rotavirus NSP3 is a translational surrogate of the poly(A)-binding protein-poly(A) complex. J Virol 89: 8773–8782.

28. Piron M, Delaunay T, Grosclaude J, Poncet D. 1999. Identification of the RNA-binding, dimerization, and eIF4GI-binding domains of rotavirus nonstructural protein NSP3. J Virol 73: 5411–5421.

29. Duan L, Zheng Q, Zhang H, Niu Y, Lou Y, Wang H. 2020. The SARS-CoV-2 spike glycoprotein biosynthesis, structure, function, and antigenicity: implications for the design of spike-based vaccine immunogens. Front Immunol 11: 576622.

30. Huang Y, Yang C, Xu XF, Xu W, Liu SW. 2020. Structural and functional properties of SARS-CoV-2 spike protein: potential antivirus drug development for COVID-19. Acta Pharmacol Sin 41 (9): 1141–1149.

31. Medina-Enríquez MM, Lopez-León S, Carlos-Escalante JA, Aponte-Torres Z, Cuapio A, Wegman-Ostrosky T. 2020. ACE2: the molecular doorway to SARS-CoV-2. Cell Biosci 10 (1): 148.

32. Rogers TF, Zhao F, Huang D, Beutler N, Burns A, He WT, Limbo O, Smith C, Song G, Woehl J, Yang L, Abbott RK, Callaghan S, Garcia E, Hurtado J, Parren M, Peng L, Ramirez S, Ricketts J, Ricciardi MJ, Rawlings SA, Wu NC, Yuan M, Smith DM, Nemazee D, Teijaro JR, Voss JE, Wilson IA, Andrabi R, Briney B, Landais E, Sok D, Jardine JG, Burton DR. 2020. Isolation of potent SARS-CoV-2 neutralizing antibodies and protection from disease in a small animal model. Science 369 (6506): 956–963.

33. Graham C, Seow J, Huettner I, Khan H, Kouphou N, Acors S, Winstone H, Pickering S, Pedro Galao R, Jose Lista M, Jimenez-Guardeno JM, Laing AG, Wu Y, Joseph M, Muir L, Ng WM, Duyvesteyn HME, Zhao Y, Bowden TA, Shankar-Hari M, Rosa A, Cherepanov P, McCoy LE, Hayday AC, Neil SJD, Malim MH, Doores KJ. 2021. Impact of the B.1.1.7 variant on neutralizing monoclonal antibodies recognizing diverse epitopes on SARS-CoV-2 Spike. bioRxiv [Preprint]. 2021 Feb 3:2021.02.03.429355.

34. Xiaojie S, Yu L, Lei Y, Guang Y, Min Q. 2021. Neutralizing antibodies targeting SARS-CoV-2 spike protein. Stem cell research 50: 102125.

35. Liu L, Wang P, Nair MS, Yu J, Rapp M, Wang Q, Luo Y, Chan JF, Sahi V, Figueroa A, Guo XV, Cerutti G, Bimela J, Gorman J, Zhou T, Chen Z, Yuen KY, Kwong PD, Sodroski JG, Yin MT, Sheng Z, Huang Y, Shapiro L, Ho DD. 2020. Potent neutralizing antibodies against multiple epitopes on SARS-CoV-2 spike. Nature 584 (7821): 450–456.

36. Arnold M, Patton JT, McDonald SM. 2009. Culturing, storage, and quantification of rotaviruses. Curr Protoc Microbiol Chapter 15: Unit 15C.3.

37. Desselberger U. 2020. What are the limits of the packaging capacity for genomic RNA in the cores of rotaviruses and of other members of the Reoviridae? Virus Res. 276:197822.

38. Meyer B, Chiaravalli J, Gellenoncourt S, Brownridge P, Bryne DP, Daly LA, Walter M, Agou F, Chakrabarti LA, Craik CS, Eyers CE, Eyers PA, Gambin Y, Sierecki E, Verdin E, Vignuzzi M, Emmott E. 2020. Characterisation of protease activity during SARS-CoV-2 infection identifies novel viral cleavage sites and cellular targets for drug repurposing. bioRxiv 2020.09.16.297945.

39. Roulston C, Luke GA, de Felipe P, Ruan L, Cope J, Nicholson J, Sukhodub A, Tilsner J, Ryan MD. 2016. ’2A-like’ signal sequences mediating translational recoding: a novel form of dual protein targeting. Traffic 17 (8): 923–39.

40. Luke G, Escuin H, De Felipe P, Ryan M. 2010. 2A to the fore - research, technology and applications. Biotechnol Genet Eng Rev 26: 223–60.

41. Liu Z, Chen O, Wall J, Zheng M, Zhou Y, Wang L, Vaseghi HR, Qian L, Liu J. 2017. Systematic comparison of 2A peptides for cloning multi-genes in a polycistronic vector. Scientific reports 7 (1): 2193.

42. Sharma P, Yan F, Doronina VA, Escuin-Ordinas H, Ryan MD, Brown JD. 2012. 2A peptides provide distinct solutions to driving stop-carry on translational recoding. Nucleic Acids Res. 40 (7): 3143–51.

43. Shaimardanova AA, Kitaeva KV, Abdrakhmanova II, Chernov VM, Rutland CS, Rizvanov AA, Chulpanova DS, Solovyeva VV. 2019. Production and application of multicistronic constructs for various human disease therapies. Pharmaceutics 11 (11): 580.

44. Philip AA, Dai J, Katen SP, Patton JT. 2020. Simplified reverse genetics method to recover recombinant rotaviruses expressing reporter proteins. J Vis Exp 158: e61039.

45. Matthijnssens J, Ciarlet M, McDonald SM, Attoui H, Bányai K, Brister JR, Buesa J, Esona, MD, Estes MK, Gentsch JR, Iturriza-Gómara M, Johne R, Kirkwood CD, Martella V, Mertens PP, Nakagomi O, Parreño V, Rahman M, Ruggeri FM, Saif LJ, Santos N, Steyer A., Taniguchi K, Patton JT, Desselberger U, Van Ranst M. 2011. Uniformity of rotavirus strain nomenclature proposed by the Rotavirus Classification Working Group (RCWG). Arch Virol 156: 1397–1413.

